# Global inhibition of deadenylation stabilizes the transcriptome in mitotic cells

**DOI:** 10.1101/2025.07.22.666109

**Authors:** Ekaterina Khalizeva, Arash Latifkar, David P. Bartel, Jimmy Ly, Iain M. Cheeseman

## Abstract

In the presence of cell division errors, mammalian cells can pause in mitosis for tens of hours with little to no transcription, while still requiring continued translation for viability. These unique aspects of mitosis require substantial adaptations to the core gene expression programs. Indeed, during interphase, the homeostatic control of mRNA levels involves a constant balance of transcription and degradation, with a median mRNA half-life of ∼2–4 hours. If such short mRNA half-lives persisted in mitosis, cells would be expected to quickly deplete their transcriptome in the absence of new transcription. Here, we report that the transcriptome is globally stabilized during prolonged mitotic delays. Typical mRNA half-lives are increased >4-fold in mitosis compared to interphase, thereby buffering mRNA levels in the absence of new synthesis. Moreover, the poly(A)-tail-length profile of mRNAs changes in mitosis, strongly suggesting a mitotic repression of deadenylation. We further show that mRNA stabilization in mitosis is dependent on cytoplasmic poly(A)-binding proteins PABPC1&4. Depletion of PABPC1&4 disrupts the maintenance of mitotic arrest, highlighting the critical physiological role of mitotic transcriptome buffering.

**Highlights:**

- The cellular transcriptome is globally stabilized during prolonged mitotic arrest
- Distinct poly(A)-tail-length profile of mRNAs in mitosis suggests repression of deadenylation
- mRNA stabilization in mitosis is dependent on PABPC1 and PABPC4
- Degradation of mRNAs during mitosis compromises maintenance of mitotic arrest

## Introduction

Cellular mRNA levels are controlled by constant turnover, balancing transcription and mRNA degradation. This homeostatic control promotes remarkable resilience in the presence of various physiological stressors, allowing cells to compensate for perturbations in mRNA synthesis by reducing degradation, and vice versa (Hartenian & Glaunsinger, 2019; Slobodin *et al*, 2020; Sun *et al*, 2013; Haimovich *et al*, 2013). The global regulation of mRNA stability thus becomes particularly important in contexts in which mammalian cells attenuate transcription for prolonged periods (Hartenian & Glaunsinger, 2019; Helenius *et al*, 2011). For example, mouse oocytes have very low levels of RNA synthesis during meiotic maturation, which takes weeks to occur (Brower *et al*, 1981; Jahn *et al*, 1976; Moore & Lintern-Moore, 1978). Oocytes resume transcription only after fertilization and zygotic genome activation. During these transcriptionally silent stages of oogenesis and early embryogenesis, maternal mRNA stability is mediated, at least in part, by the germline-specific RNA-binding protein MSY29 (Medvedev *et al*, 2011) and by subcellular mRNA localization (Cheng *et al*, 2022). The maintenance of maternal mRNAs is crucial for timely translational activation and proper oocyte maturation during and after the transcriptionally silent stages of development (Conti & Kunitomi, 2024; Winata & Korzh, 2018).

Mitotic cells also exhibit significantly downregulated transcription, as evidenced by chromosome condensation (Antonin & Neumann, 2016), eviction of transcriptional machinery from chromatin (Parsons & Spencer, 1997; Martínez-Balbás *et al*, 1995; Perea-Resa *et al*, 2020), inactivation of general transcription factors (Segil *et al*, 1996), and low levels of nascent transcript labeling (Taylor, 1960; Palozola *et al*, 2017). Although mitosis typically lasts for ∼1 hour, errors induced by either physiological damage or anti-mitotic drugs can cause somatic cells to remain arrested in mitosis for hours (Gascoigne & Taylor, 2008; Skoufias *et al*, 2006). This arrest is mediated by the conserved Spindle Assembly Checkpoint signaling pathway (McAinsh & Kops, 2023). During an extended mitotic delay, ongoing translation is essential for cell survival and the ability to maintain a cell cycle arrest (Tsang & Cheeseman, 2023; Ly *et al*, 2024). Following the resolution of any mitotic damage, cells can segregate their chromosomes, exit mitosis, and re-enter G1 (Sinha *et al*, 2019), once again becoming transcriptionally competent (Palozola *et al*, 2017).

In asynchronous mammalian somatic cells, the median mRNA half-life (t_1/2_) is typically 2–4 h, depending on the cell type and the half-life measurement strategy (Eisen *et al*, 2020; Herzog *et al*, 2017; Lugowski *et al*, 2018). This suggests that, during prolonged periods of transcriptional silencing in which ongoing translation is essential, cells may require compensatory mechanisms to prevent depletion of their transcriptomes. Interestingly, somatic cells do not express the factors that promote mRNA stabilization in oocytes, such as MSY29, which suggests that there must be a distinct strategy for regulating mRNA stability. Here, we report that mitotic cells globally stabilize their transcriptomes to maintain an extended mitotic arrest in the near-complete absence of mRNA synthesis. We find that the poly(A)-tail-length profile of mRNAs in mitotic cells is indicative of repressed deadenylation. In addition, we find that cytoplasmic poly(A)-binding proteins (PABPC1&4) are necessary for mitotic mRNA stabilization. Induction of mRNA degradation in mitosis through the depletion of PABPC1&4 disrupts the cells’ ability to maintain prolonged mitotic arrest, highlighting the physiological importance of buffering transcriptome levels in mitosis.

## Results

### The cellular transcriptome is not depleted during a prolonged mitotic arrest

To test whether mitotically arrested cells maintain their transcriptome levels despite the prolonged global inhibition of transcription, we first evaluated changes in mRNA levels during an extended mitotic arrest. We synchronized HeLa cells using a double thymidine block followed by treatment with the KIF11 inhibitor S-trityl-L-cysteine (STLC). STLC triggers a mitotic arrest (Fig. EV1A) by activating the Spindle Assembly Checkpoint but does not activate stress response pathways (Ly *et al*, 2024; Skoufias *et al*, 2006; Musacchio, 2015). We then performed RNA-seq at 0–1 h, 6 h, and 24 h after entry into mitosis. Normalization to spike-in RNA allowed us to quantitatively compare global RNA level changes between different timepoints (Fig. 1A). Strikingly, the abundances of most mRNAs remained very close to their initial levels even after 24 hours of mitotic arrest (Fig. 1B–D; Fig. EV1B). The maintenance of transcriptome levels was not due to a resumption of transcription in the arrested cells, as FACS-based analysis to detect the incorporation of the uridine analogue 5-Ethynyluridine (5EU) at the relevant timepoints confirmed a lack of ongoing transcription (Fig. 1E). Together, these results indicate that cells maintain transcriptome levels throughout a 24-hour mitotic arrest, despite a near-complete inhibition of mRNA synthesis, suggesting that a mechanism must be in place to prevent global mRNA depletion.

**Figure 1.**
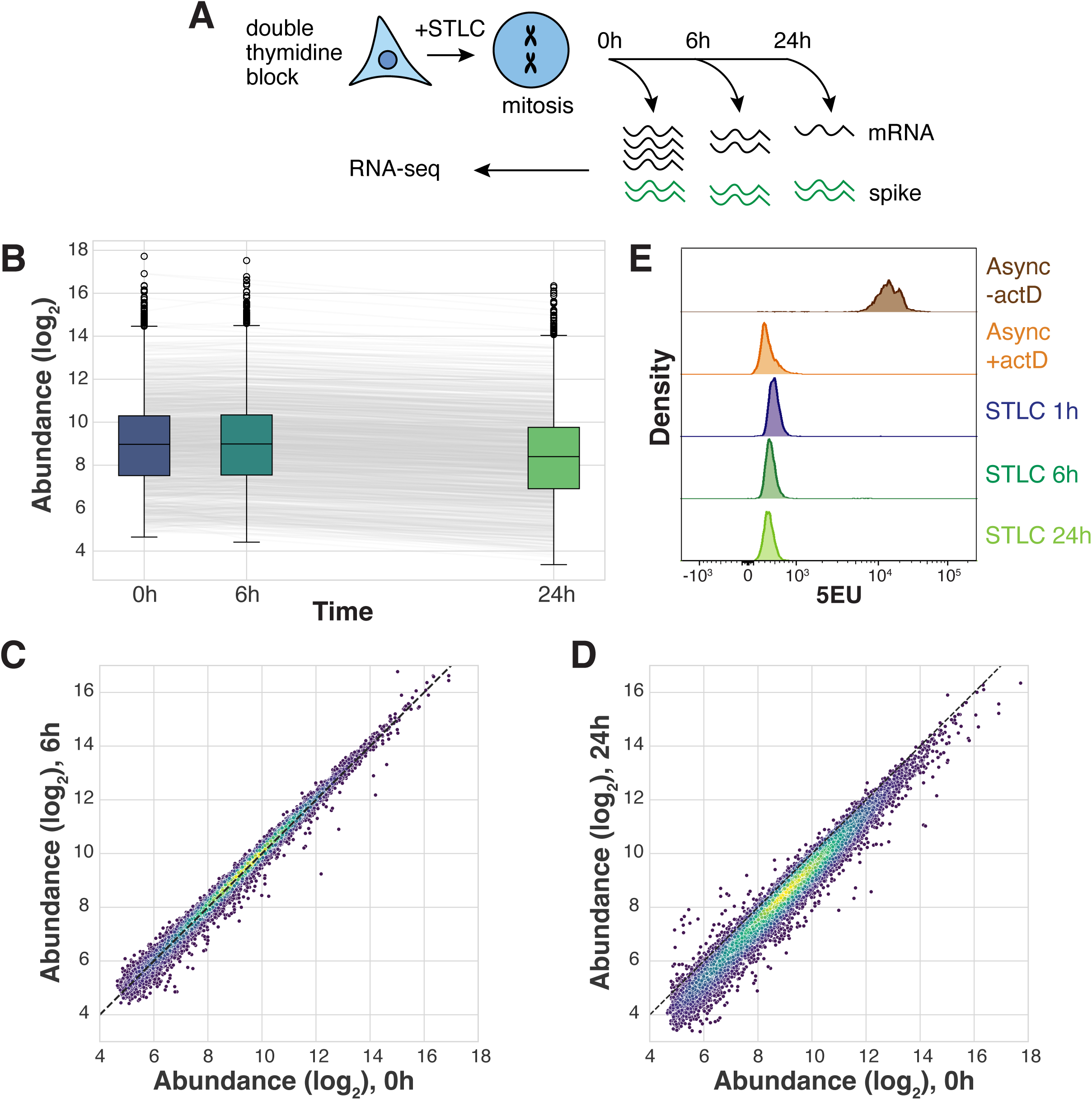
Cellular transcriptome does not get depleted during prolonged mitotic arrest. **A)** Schematic diagram of timecourse experiments to measure mRNA levels during STLC-induced mitotic arrest. **B)** Boxplots comparing spike-normalized mRNA abundances over the course of 24 h of mitotic arrest. Each line represents the normalized abundance of an mRNA across the timecourse. Each point represents the mean of two biological replicates, n = 11282. **C-D**) Scatterplots comparing spike-normalized abundances between (**C**) 0 h and 6 h or (**D**) 0 h and 24 h of arrest. Dotted line indicates x=y diagonal, points colored by density. Each point represents the mean of two biological replicates, n = 11282. **E**) Histogram comparing 5EU incorporation in asynchronous samples with or without actinomycin D, and in STLC-arrested samples.

### mRNA half-lives are increased in mitotic cells

To test whether mRNA decay is perturbed in mitotically arrested cells, we next measured mRNA levels over time following pharmacological inhibition of transcription by actinomycin D in cells arrested in either interphase (G2 stage) or M phase of the cell cycle using the CDK1 inhibitor RO-3306 or STLC, respectively (Fig. 2A, Fig. EV2A). We then calculated mRNA half-lives by fitting abundance values to an exponential curve (Fig. EV2B; Methods). In G2 cells, we observed a median mRNA half-life of 5.6 h, in range with prior measurements of mRNA stability in actinomycin-D-treated interphase cells (Herzog *et al*, 2017; Lugowski *et al*, 2018). However, mRNA half-lives were almost 4-fold higher in mitotically arrested cells, with a median half-life of 20.1 h (Fig. 2B). Notable examples of mRNA stabilization include transcripts such as *MYC*, *KLF10*, and *ATF3*, which are unstable in G2 (t_1/2_ 1.0 h, 1.4 h, and 1.9 h, respectively) but underwent little to no apparent degradation in mitotic cells (Fig. 2C). An even stronger global stabilization was observed when comparing t_1/2_ in G1 vs. M cell cycle stages (median t_1/2_ of 3.1 h in G1 vs. 27.3 h in M; Fig. 2D). This result is in line with previously reported waves of mRNA decay upon M-to-G1 transition (Krenning *et al*, 2022). We also observed a similar effect when using the CDK7 inhibitor THZ1 instead of actinomycin D to inhibit RNA polymerase II-dependent transcription (median t_1/2_ of 6.8 h in G2 vs. 33.4 h in M; Fig. EV2C).

**Figure 2.**
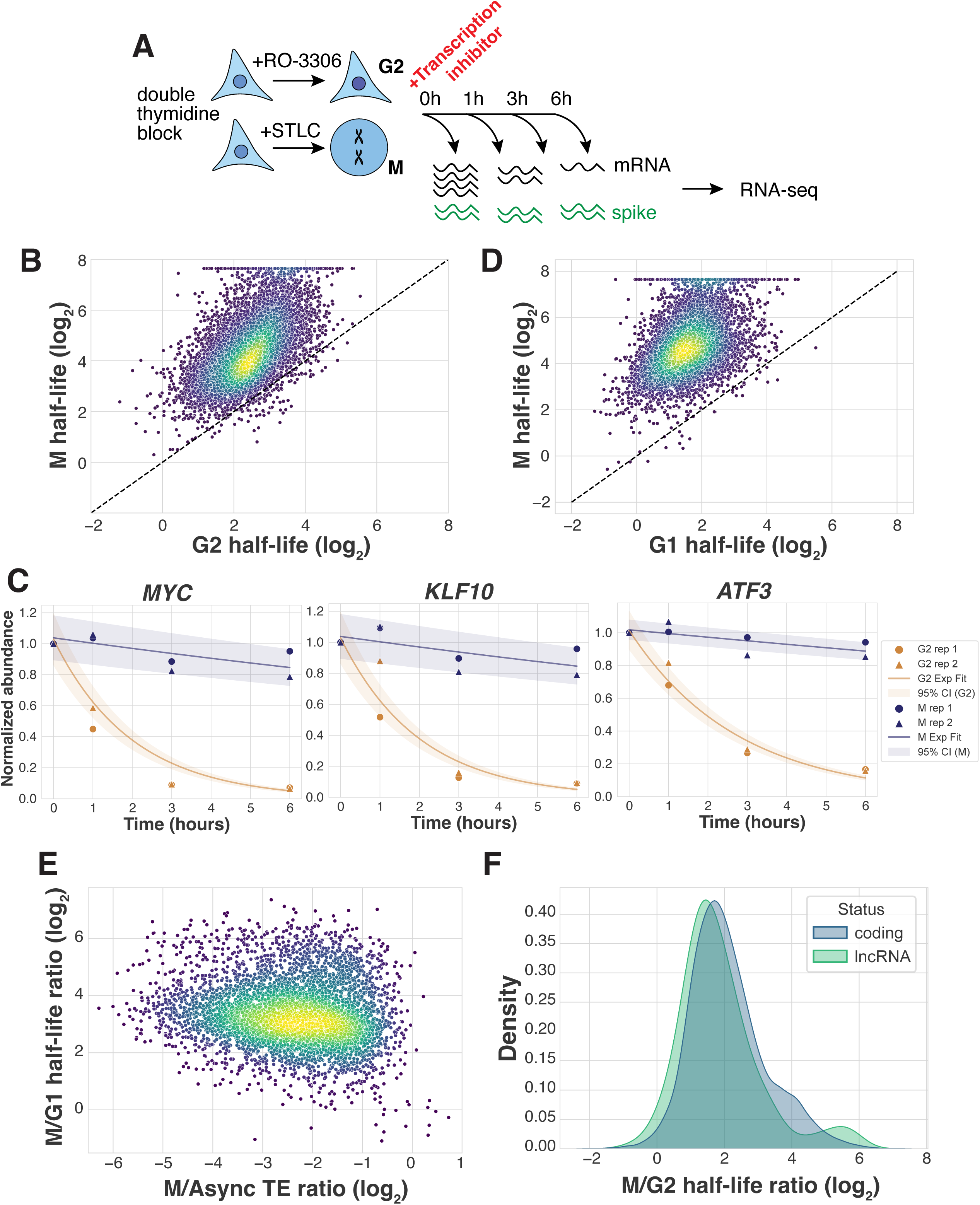
mRNA half-lives are increased in cells arrested in mitosis compared to interphase. **A)** Schematic diagram of transcriptional inhibition timecourse experiments to measure mRNA half-lives in G2- and M-synchronized cells. **B)** Scatterplot comparing mRNA half-lives in G2- and M-arrested cells treated with actinomycin D. Each point represents the mean of 2 biological replicates, n = 9858. Median t_1/2_ 5.58 h and 20.13 h for G2 and M, respectively. **C)** Exponential curve fit to individual transcript abundances, plotted with 95% confidence intervals. *MYC* t_1/2_ 1.02 h and 21.19 h, *R*^2^ 0.99 and 0.65 in G2 and M, respectively. *KLF10* t_1/2_ 1.4 h and 19.47 h, *R*^2^ 0.97 and 0.55 in G2 and M, respectively. *ATF3* t_1/2_ 1.91 h and 29.01 h, *R*^2^ 0.98 and 0.74 in G2 and M, respectively. **D)** Scatterplot comparing mRNA half-lives in cells synchronized in G1 or M treated with actinomycin D. Data from one biological replicate, n = 6969. Median t_1/2_ 3.06 h and 27.3 h for G1 and M, respectively. **E)** Scatterplot comparing transcript stabilization (M/G1 half-life ratio) and changes in translation efficiency (M/Async translation efficiency ratio, data from (Ly *et al*, 2024)). n = 4772, *R*_s_ = –0.12. **F)** Probability density function plot comparing degree of transcript stabilization (M/G2 half-life ratio) for coding transcripts and lncRNAs. Median stabilization 3.82-fold for coding transcripts and 3.08-fold for lncRNAs.

Prior work identified mRNA properties that are associated with altered RNA stability, such as AU-rich elements (Chen & Shyu, 2011), P-body localization (Blake *et al*, 2024), and ORF exon density (Spies *et al*, 2013; Agarwal & Kelley, 2022). However, none of these tested features showed a strong correlation with the degree of mitotic stabilization (Fig. EV2D–G). These observations are consistent with a global reduction in mRNA degradation that is not dependent on targeted mRNA destabilization factors.

Prior work suggested that there is feedback between translation and mRNA stability in asynchronously growing cells (Jia *et al*, 2020). Despite a requirement for ongoing protein production, translation is partially attenuated in mitotic cells (Tanenbaum *et al*, 2015; Ly *et al*, 2024). Therefore, we considered whether the observed global transcriptome stabilization during mitosis could be explained by the global decrease in translation (Morris *et al*, 2021; Rosa-Mercado *et al*, 2024). To test this, we evaluated the degree of stabilization of each transcript relative to its change in translation efficiency between asynchronous and mitotic samples, as measured by ribosome profiling (Ly *et al*, 2024). This analysis detected no correlation between the changes in mRNA half-life and translation efficiency (Fig. 2E, *R_s_*= –0.12). Notably, mRNAs that lacked translational changes during mitosis were still stabilized. We also compared the measured change in stability with the translation status of a given transcript. Long non-coding RNAs (lncRNAs), which typically do not undergo translation, displayed only a modestly lower degree of stabilization in mitosis compared to coding transcripts (3.1- and 3.8-fold changes, respectively; Fig. 2F). Together, these results suggest that there is a striking transcriptome-wide mRNA stabilization during mitosis, and that a global attenuation of translation cannot explain this behavior.

### Deadenylation is globally perturbed during mitosis

The global increase in mRNA half-lives that we observed in mitotic cells suggests that mRNA decay is perturbed. mRNA degradation is a multi-step process that involves sequential reactions (Garneau *et al*, 2007; Wiederhold & Passmore, 2010; Passmore & Coller, 2022; Dowdle & Lykke-Andersen, 2025). For most mRNAs, decay begins with shortening of the poly(A)-tail by cytoplasmic deadenylases (Eisen *et al*, 2020; Raisch & Valkov, 2022). A subset of deadynylases can displace the poly(A)-binding protein (PABP) and gradually digest the poly(A) tail until it is too short to be bound by a single PABP (<20 nt) (Yi *et al*, 2018; Webster *et al*, 2018). After deadenylation, decapping enzymes are recruited to remove the m7G cap that stabilizes the 5’ end of the mRNA (Van Dijk, 2002). The body of the transcript is then subject to digestion by 5’-to-3’ and 3’-to-5’ exonucleases (Chang *et al*, 2014; Anderson, 1998). Inhibition of any of the steps and their corresponding enzymes during mitosis could result in global transcriptome stabilization. As the first and rate-limiting step of the mRNA degradation pathway is deadenylation (Chen & Shyu, 2011; Eisen *et al*, 2020; Passmore & Coller, 2022), we chose to start by analyzing changes in poly(A) tail length.

To evaluate tail-length distributions, we conducted poly(A)-tail-length profiling by sequencing (PAL-seq) (Subtelny *et al*, 2014) in cells arrested in mitosis or G2 after up to 6 h of transcriptional inhibition (Fig. 3A). In G2-arrested cells, the initial tail-length distribution was consistent with previous reports of the bulk poly(A) tail length profile in asynchronous cells (Subtelny *et al*, 2014), with a peak at around 40–50 nt and a strong right skew (Fig. 3B). In the absence of new mRNA synthesis, which typically acts to replenish long-tailed species lost in ongoing deadenylation, the bulk poly(A) tail length distribution in G2 cells shifted leftward over time, reflecting a progressive depletion of longer-tailed species and overall tail length shortening. The shift was particularly noticeable in the 0–30 nt and >150 nt ranges (Fig. 3B), but was modest overall, presumably because many of the short-tailed species generated over the time course had been degraded.

**Figure 3.**
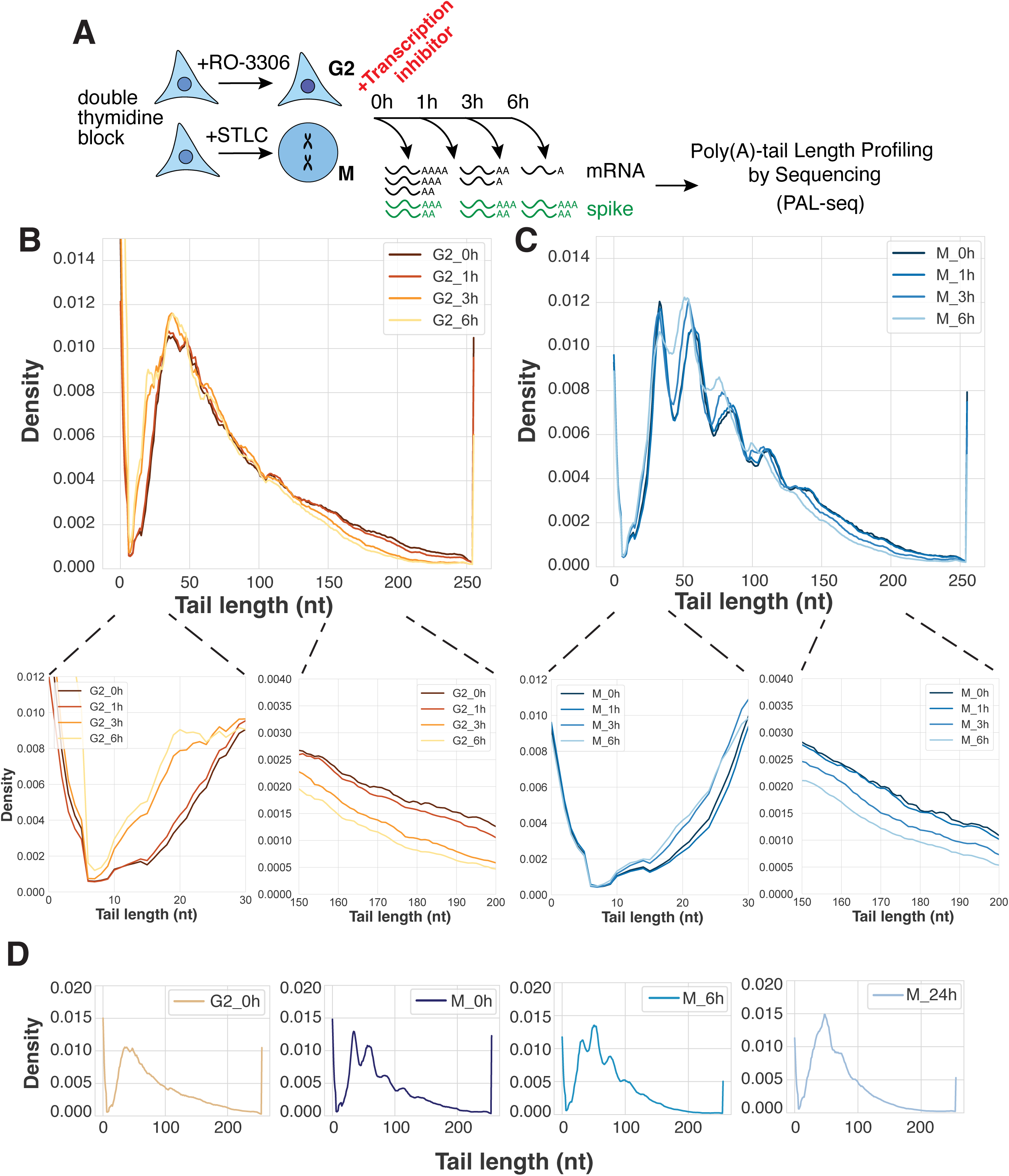
Deadenylation is globally perturbed during mitotic arrest. **A)** Schematic diagram of transcriptional inhibition timecourse experiments to measure poly(A) tail length profiles in synchronized cells by PAL-seq. **B–C**) Poly(A) tail length distributions in (**C**) G2- or (**D**) M-arrested cells treated with actinomycin D as measured by PAL-seq. Insets show distribution between 0–30 nt and 150–200 nt. **D**) Poly(A) tail length distributions in M-arrested cells without transcriptional inhibitors as measured by PAL-seq. A trace from G2-arrested cells prior to actinomycin D treatment (shown in **B**) is included for reference.

Intriguingly, the tail length distributions in mitotic cells appeared markedly different from those of G2-arrested cells, suggesting repressed deadenylation during mitosis (Fig. 3C). At the initial timepoint, the median poly(A) tail length in G2 cells was 72 nt, compared to 76 nt in mitotic cells. After 6 h of transcriptional inhibition, the median tail length in G2 cells decreased to 58 nt, compared to 67 nt in mitotic cells, exhibiting a 14 nt and 9 nt shortening, respectively. Mitotic distributions still showed shortening of long tails in the >150 nt range after transcriptional inhibition. However, the shift in the 0– 30 nt range became notably less prominent, suggesting that shorter-tailed transcripts are not deadenylated as efficiently during mitosis as they are in interphase. In principle, another possibility is that short-tailed isoforms are more efficiently cleared in mitotic cells. However, this alternative possibility is not consistent with the mRNA stabilization we observed during mitosis (Fig. 1B–D, 2B–D).

In addition to the reduction in overall poly(A) tail-length shortening, we observed distinct “bumps” in the mitotic tail-length distributions, occurring with a ∼30 nt periodicity as early as 0–1 h after entry into mitosis and persisting throughout a 6 h mitotic arrest (Fig. 3C). These bumps were notably absent from the spiked-in zebrafish poly(A) tails (Fig. EV3A). A similar shape of the tail-length distribution is reported in conditions of >3-hour transcription inhibition in mouse cells (Eisen *et al*, 2020) or following inactivation of the CCR4–NOT complex (Yi *et al*, 2018). When considering that the cytoplasmic poly(A)-binding protein (PABPC) covers about 27 nt of the poly(A) tail (Baer & Kornberg, 1980), this distinctive patterning is attributed to more dense, and therefore more phased, packing of PABPC on the poly(A) tail, coupled with reduced CCR4–NOT 3’ exonuclease activity upon reaching a PABPC-protected region of the tail (Yi *et al*, 2018; Eisen *et al*, 2020). We will therefore refer to the observed periodic bumps as “phased PABPC toeprints” (noting that toeprints, in contrast to “footprints,” mark only one end of a binding site (Nou & Kadner, 2000)). Importantly, the trends that we observed in the transcriptome-wide data were also apparent at the level of individual transcripts (Fig. EV3B–E). Analysis of the 25% most highly expressed transcripts and the 25% lowest expressed transcripts confirmed the reduced leftward shift in the low tail range and the similar pattern of phased toeprints in each case (Fig. EV3F–G). We observed a similar behavior regardless of the degree of mitotic mRNA stabilization, with the 25% most stabilized and 25% least stabilized transcripts exhibiting the same trends (Fig. EV3H–I). Thus, the prominent periodic PABPC toeprints in mitotic samples are not driven solely by a few highly expressed genes, and do not correlate with the degree of mitotic transcript stabilization.

To assess whether deadenylation is repressed during a prolonged mitotic arrest in the absence of pharmacological transcription inhibition, we conducted PAL-seq on cells arrested in mitosis over the course of 24 hours. Tail-length distributions in these cells were consistent with those measured in transcriptionally inhibited cells (Fig. 3D). We observed the same pattern of phased PABPC toeprints as in actinomycin-D-treated cells, with well-defined bumps appearing as early as 0–1 h into mitosis and persisting for hours.

Together, these observations suggest that deadenylation is perturbed in mitotically arrested cells, leading to inefficient tail length shortening in the absence of new mRNA synthesis. The striking phasing of PABPC toeprints, which becomes apparent almost immediately as cells enter mitosis, suggest that PABPC is not efficiently removed from the poly(A) tails in this cell cycle stage. These data support a model in which deadenylation is perturbed in mitotically arrested cells, resulting in a global decrease in mRNA degradation and allowing cells to maintain their transcriptomes in the near absence of new mRNA synthesis.

### *UBE2C* escapes transcriptome-wide mitotic stabilization

The change in mRNA stability and deadenylation occurs globally and affects the vast majority of the analyzed transcripts. However, we identified *UBE2C*, a transcript encoding an E2 ubiquitin ligase, as a notable exception to the transcriptome-wide trends (Fig. EV4). UBE2C functions together with the anaphase promoting complex/cyclosome (APC/C), a cell cycle-regulated E3 ubiquitin ligase that controls progression through mitosis by targeting various substrates for proteasomal degradation (Jin *et al*, 2008). Interestingly, in our analysis, the *UBE2C* mRNA consistently behaved contrary to the transcriptome-wide trends. In samples arrested in mitosis by STLC treatment for 24 hours, *UBE2C* displayed rapid degradation, with a fitted t_1/2_ of 2.1 h (Fig. EV4A). Similarly, in transcriptionally arrested G2- and M-synchronized cells, *UBE2C* was destabilized in mitosis compared to interphase (t_1/2_ of 1.8 h in M and 6.3 h in G2; Fig. EV4B). Perhaps most strikingly, the distributions of poly(A) tail lengths for this transcript were also reversed in interphase and mitosis, with prominent phasing in G2 samples and no phasing in mitotic samples (Fig. EV4C). Therefore, our analysis suggests that some transcripts may have evolved ways to evade transcriptome-wide stabilization in mitosis.

### PABPC1&4 are required for mitotic transcriptome stabilization

In interphase cells, deadenylases efficiently remove PABPC from poly(A) tails to allow for tail shortening. The presence of prominent periodic toeprints prompted us to hypothesize that PABPC stabilizes transcripts in mitosis by protecting poly(A) tails from deadenylation and consequent degradation. To assess this possibility, we used a PABPC1&4-Auxin-Inducible Degron (AID) system (Xiang & Bartel, 2021) to evaluate the effects of rapid PABPC1&4 depletion on mRNA stability in mitotic cells. To selectively eliminate PABPC1&4 in mitosis without affecting prior interphase behaviors, we synchronized cells in mitosis using STLC treatment, after which IAA was added to induce rapid OsTIR1-mediated degradation of PABPC1&4 (Fig. 4A–B). In control cells expressing PABPC1&4, mRNA levels did not change substantially after 4 hours of mitotic arrest, consistent with our original finding (Fig. 4C, D). Strikingly, in PABPC1&4-depleted cells, the levels of most mRNAs decreased within 4 hours of IAA addition, suggesting that the mRNA degradation pathway is active in mitotic cells in the absence of PABP. Notably, the range of fold-change values was also wider in PABP-depleted mitotic cells than in the control condition, consistent with restored regulation of mRNA stability (Fig. 4D). This result suggests that the global transcriptome stabilization observed in mitotically arrested cells occurs through the inhibition of deadenylation.

**Figure 4.**
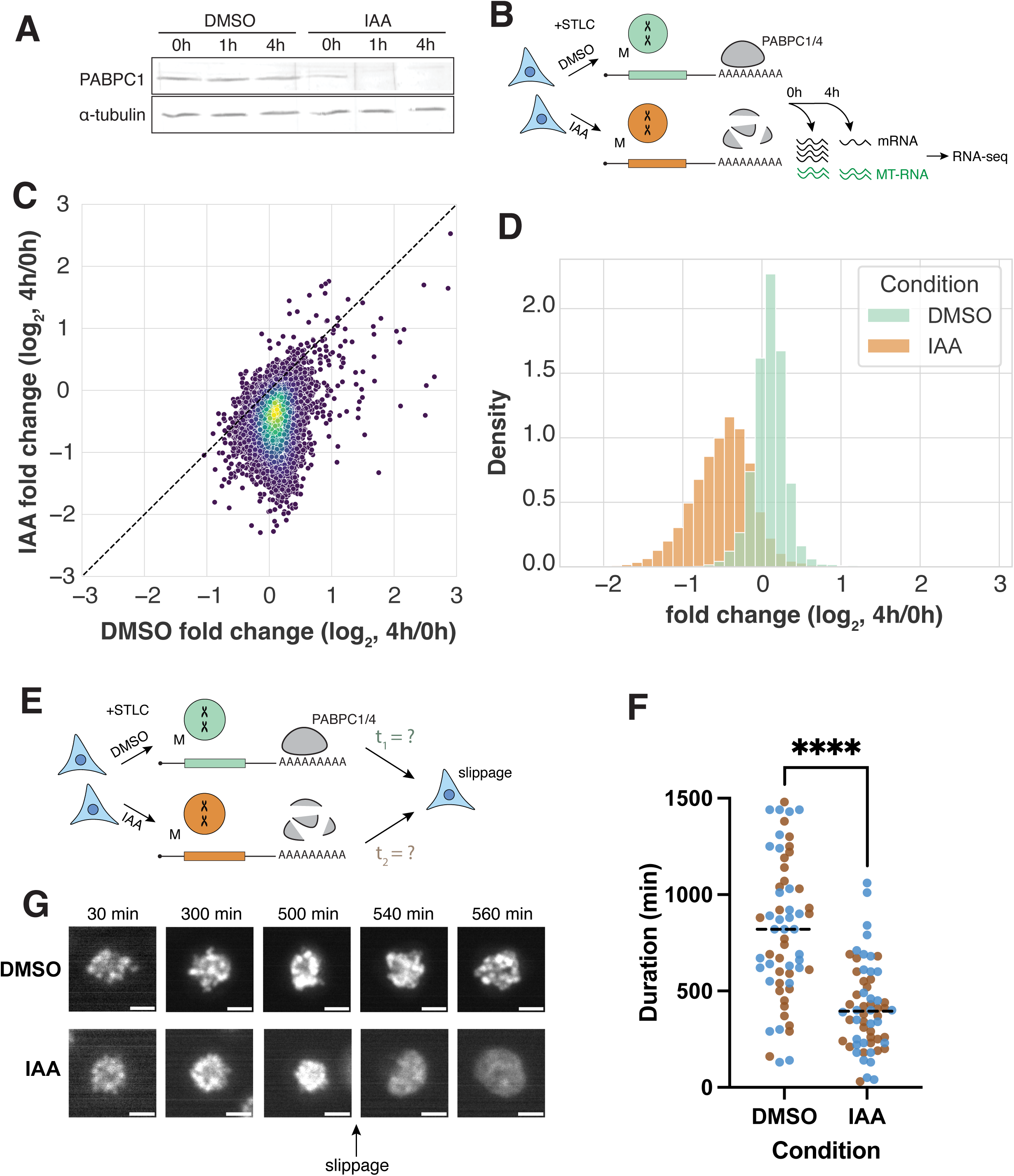
PABPC1&4 are required for mitotic transcriptome stabilization and maintenance of mitotic arrest. **A)** Western blot showing PABPC1 depletion in STLC-arrested cells following either DMSO treatment, or IAA-induced degradation of PABPC1&4. alpha-tubulin shown as a loading control. **B)** Schematic diagram of sequencing experiments with PABPC1&4 depletion in M-arrested cells. **C)** Scatterplot comparing mRNA abundance fold changes in M-arrested HCT116 cells within 4 hours of either DMSO control treatment or IAA-induced PABPC1&4 depletion. Each point represents the mean of two replicates, n = 11078. Median fold changes: 1.07 and 0.72 for control and PABPC1&4 depletion, respectively. **D)** Probability density function plot comparing expression fold-change within 4 hours of either DMSO control treatment or IAA-induced PABPC1&4 degradation. **E)** Schematic diagram of live imaging experiments with PABPC1&4 depletion to measure mitotic slippage in STLC-arrested cells. **F)** Dot plot comparing time from control treatment or PABPC1&4 degradation until mitotic slippage in STLC-treated HCT116 cells. n = 30 cells per condition per replicate, points colored by replicate. Median durations: 820 min for DMSO control cells and 395 min for PABPC1&4-depleted cells. Student’s *t*-test *p* < 0.0001. **G)** Representative images of SiR-DNA-stained DNA in STLC-arrested HCT116 cells with (DMSO) or without (IAA) PABPC1&4. Brightness not scaled identically to highlight differences between condensed and de-condensed chromosomes.

### mRNA stabilization is necessary for maintenance of mitotic arrest

Cells arrested in mitosis using STLC treatment or other mitotic disruptions can remain viable for tens of hours despite the near absence of transcription. In mitotically arrested cells, ongoing translation is necessary for the maintenance of mitotic arrest (Malureanu *et al*, 2010; Mena *et al*, 2010; Tsang & Cheeseman, 2023). Therefore, we hypothesized that mitotic transcriptome stabilization would also be required for cells to continue translation of key mitotic factors and thus persist in the absence of new transcription during mitotic arrest. To assess the physiological importance of mitotic transcriptome stabilization, we tracked the fate of STLC-treated HCT116 cells in the presence or absence of PABPC1&4 using the AID system (Fig. 4E).

Cells challenged with STLC face two mutually exclusive fates: they either die during mitosis or exit mitosis without chromosome segregation, entering G1 through a process known as “mitotic slippage” (Skoufias *et al*, 2006). In contrast to control cells that remained arrested in mitosis for hours, PABPC1&4 depletion resulted in >2-fold accelerated mitotic slippage (Fig. 4F, median times of 820 min for control vs 395 min for PABP-depleted cells). Thus, in the absence of transcriptome stabilization, cells are unable to maintain a mitotic arrest to fix cell division errors and instead “slip out” into a tetraploid G1 state without dividing.

Together, these results suggest that repression of deadenylation in mitotic cells is important for globally preserving mRNA levels during an extended mitotic arrest. Crucially, if cells cannot stabilize mRNA in mitosis, their ability to maintain a prolonged arrest is significantly impaired, suggesting that this mechanism is critical for proper mitotic physiology.

## Discussion

Our study uncovers a previously unappreciated phenomenon in which mammalian cells can maintain a state of mitotic arrest for tens of hours without global transcriptome changes. We demonstrate that mRNA expression levels are maintained over a 24-hour mitotic arrest period in the near absence of transcription. The transcriptome is preserved due to a global increase in mRNA stability in mitotic cells. This mRNA stabilization is mediated by the attenuation of deadenylation, as evident from slowed shortening of poly(A) tail lengths, characteristic phasing of PABPC-protected toeprints, and the dependence of this stabilization on the presence of PABPC1&4.

Prior reports suggested that there is a transcriptome buffering feedback, with cells in which transcription is inhibited for 24 hours adapting to this change by reducing mRNA decay (Slobodin *et al*, 2020). Although such a feedback mechanism could be partially responsible for the stabilization in mitotically arrested cells observed in this study, this is inconsistent with the rapid changes we observed in both mRNA half-lives and poly(A)-tail-length profiles upon entry into mitosis. For instance, in prior studies of transcription-degradation feedback, the effect of transcription perturbation on poly(A)-tail profiles did not occur until >3 hours after drug treatment in NIH 3T3 cells (Eisen *et al*, 2020). In contrast, in mitotically arrested cells, the characteristic phased PABPC toeprints become clear as early as 0–1 h after entry into mitosis (Fig. 3). Additionally, if transcriptional inhibition was solely responsible for mRNA stabilization in mitotic cells, our experiments comparing mRNA half-lives in cells with pharmacologically inhibited transcription would have shown similar results in cells arrested in G2- or M-phase. Therefore, a mechanism other than transcription-degradation feedback must contribute to prompt stabilization of the transcriptome upon entry into mitosis.

We hypothesize that inactivation of the CCR4–NOT complex may contribute to mRNA stabilization in mitosis. Although in principle altered polyadenylation could have also contributed to differences in poly(A)-tail profiles, recent work has demonstrated that mitotic phosphorylation of PABPN1 prevents hyperadenylation (Gordon *et al*, 2024). Therefore, decreased deadenylation by CCR4–NOT is the most likely explanation for the striking poly(A) tail length distributions in mitosis. Consistent with our model, prior work has shown that inactivation of the CCR4–NOT complex can result in phased PABPC toeprints similar to the ones observed in our study (Yi *et al*, 2018). Additionally, a recent study suggested that the CCR4–NOT complex may not associate with RNA as efficiently in mitosis as it does in interphase (Fig. EV5B; Rajagopal *et al*, 2025).

Importantly, targeted depletion of PABPC1&4 during mitosis leads to a strong reduction mRNA stability and also prevents cells from maintaining a mitotic arrest. Mechanistically, the effect of PABPC1&4 depletion on mitotic arrest maintenance is likely to involve the combined effects of stabilizing hundreds of mRNAs, with specific contributions from each transcript. For example, we propose that one relevant target of PABPC-dependent stabilization is the *CCNB1* (Cyclin B1) transcript. Continued CCNB1 translation during mitosis is required for sustaining the mitotic state through cyclin-dependent kinase activity, with CCNB1 degradation serving as a key step in mitotic exit (Mena *et al*, 2010; Malureanu *et al*, 2010). We found that the *CCNB1* transcript stability is increased 6-fold in mitotic cells compared to G2. Additionally, *CCNB1* is 36% less stable in PABPC-depleted cells compared to control cells (Fig. EV5C). Although these observations do not demonstrate a causative role of *CCNB1* mRNA degradation in the early slippage of PABPC-depleted mitotic cells or preclude contributions from other mRNAs, they are consistent with the expectations based on previously established effects of CCNB1 depletion on the maintenance of the mitotic state (Mena *et al*, 2010).

Together, our findings reveal a novel gene-regulatory framework in mitotic cells that is responsible for maintaining mRNA levels required for cells to endure periods of transcriptional silencing. Importantly, our results could have therapeutic implications, as antimitotic drugs such as paclitaxel are widely used as frontline cancer therapeutics (van Vuuren *et al*, 2015). Although such drugs are effective at specifically targeting dividing cells, they are also known to cause a range of adverse side effects, highlighting a need for improvements in their efficacy (Cella *et al*, 2003; Awosika *et al*, 2025). Our study nominates PABP and the mRNA degradation machinery as potential targets to enhance the efficacy of such drugs.

## Acknowledgements

This work was supported by grants from the NIH (R35GM126930 to I.M.C.). D.P.B. is an investigator of the Howard Hughes Medical Institute. A.L. is supported by an NIH fellowship (K00CA234921). J.L. is supported in part by the Natural Sciences and Engineering Research Council of Canada. We thank all the members of the Cheeseman laboratory for helpful discussions. We thank the Whitehead Institute Genome Technology Core for the RNA sequencing, the Flow Cytometry Core for assistance with FACS experiments, and the Glass Wash core for supporting our work.

## Methods

### Tissue Culture

HeLa cells were cultured in Dulbecco’s modified Eagle medium (DMEM) supplemented with 10% heat-inactivated fetal bovine serum, 2 mM L-glutamine and 100 U ml−1 penicillin-streptomycin at 37 °C with 5% CO_2_. HCT116 cells were cultured in McCoy’s 5A (modified) medium supplemented with 10% tetracycline-free heat-inactivated fetal bovine serum, 2 mM L-glutamine and 100 U ml−1 penicillin-streptomycin at 37 °C with 5% CO_2_.

### Cell Cycle Synchronization

For experiments done with a double thymidine block, HeLa cells were grown to 20% confluency and treated with 2 mM thymidine for 18 h. Cells were released from thymidine arrest by washing twice with warm DMEM and cultured for 9 h without drugs before a second treatment with 2mM thymidine for another 14 h. Cells were released from thymidine arrest by washing twice with warm DMEM and cultured in 8 µM STLC or 6 µM RO-3306 for 8 h until ∼50% of STLC-treated cells were rounded. Mitotic cells were isolated by shake off. Actinomycin D was added to cells at a concentration of 5 ug/mL. THZ1 was added to cells at 1 uM. For G1 synchronization, cells were synchronized using the double thymidine block procedure, followed by release into 8 µM STLC for 8 h. Mitotic cells were shaken off and replated in DMEM and allowed to progress to G1 and adhere to the plate for 1 h before transcriptional inhibitor treatment. For experiments done with a single thymidine block, HeLa cells were grown to 40% confluency and treated with 2 mM thymidine for 24 h before being released into 8 µM STLC as described above. For experiments with the HCT116 PABP AID cell line, cells were grown to 20% confluency and treated with 8 µM STLC for 16 h.

### Cell Cycle Quantification with FACS

Mitotic cells were collected by shake off. Interphase cells were detached with 5mM EDTA in PBS. All cells were then washed once with PBS and spun down at 500g for 5 min. Cells were then resuspended in 500uL PBS and vortexed while 4.5mL of -20C 100% ethanol was added dropwise. After a 30-minute incubation on ice or overnight storage at 4C fixed cells were spun down at 1000 g for 5 minutes and resuspended in 500uL ethanol with 9.5mL cold PBS with 3% BSA to facilitate pelleting and prevent clumping. Cells were then washed once with 1mL PBS + 3% BSA + 0.1% Triton X-100 and incubated on ice in 1mL antibody dilution buffer (20 mM Tris-HCl, 150 mM NaCl, 0.1% Triton X-100, 3% bovine serum albumin, 0.1% NaN3, pH 7.5) for 30 minutes. Cells were then incubated in 300uL antibody dilution buffer with 1:1000 phospho-histone 3 Serine 10 antibody (Abcam 5176; Rabbit) with slow rotation overnight at 4C. After a 1mL wash in PBS + 3% BSA + 0.1% Triton X-100, cells were incubated for 1 h on ice with 300 uL antibody dilution buffer with 1:300 anti-Rabbit Cy5 antibody (Jackson ImmunoResearch Laboratories; Goat). Cells were then washed once with PBS + 3% BSA + 0.1% Triton X-100, resuspended in 500uL PBS + 20ug/mL Hoechst and incubated on ice for 30 minutes before being strained into flow cytometry tubes. Analysis of cell cycle stage was done using the BD FACSymphony A1 Cell Analyzer (BD Biosciences) and quantified using FlowJo v10.9.0. We first gated cells using forward scatter and side scatter area, followed by identifying cells with 4n DNA content and classifying them as either G2 (pH3S10-negative) or M (pH3S10-positive).

### Measurement of Global Transcription rate with FACS

HeLa cells were synchronized by double thymidine block followed by STLC treatment as described above. 0, 5, or 23 hours after the cells entered mitotic arrest, 1mM 5EU was added to cells for one hour. Asynchronous control cells received the same treatment, with the addition of 5ug/mL of actinomycin D to one of the wells. After 1 hour of 5EU labeling, mitotic cells were harvested by shake-off and asynchronous cells were dissociated from wells with 5mM PBS-EDTA. Cells were washed with 1mL cold PBS and permeabilized with 1mL of 3.7% formaldehyde in PBS for 15 min at room temperature, followed by a wash in 1mL cold 2.5% PBS-BSA. Cells were permeabilized in 1mL 0.5% Triton X-100 in PBS for 15 min at room temperature. Click chemistry was then performed using the Click-iT™ RNA Alexa Fluor™ 594 Imaging Kit (ThermoFisher Scientific, C10330) according to manufacturer’s instructions, substituting coverslip washes with 4 min 500g spins at 4C. After click chemistry, cells were resuspended in 500uL antibody dilution buffer (20 mM Tris-HCl, 150 mM NaCl, 0.1% Triton X-100, 3% bovine serum albumin, 0.1% NaN3, pH 7.5) and strained into flow cytometry tubes. Analysis of 5EU incorporation was done using the BD FACSymphony A1 Cell Analyzer (BD Biosciences) and quantified using FlowJo v10.9.0. We first gated cells using forward scatter and side scatter area, followed by plotting histograms of 5EU levels to visualize active transcription during the hour prior to cell harvesting.

### RNA Sequencing

Interphase cells were detached from plates using PBS with 5mM EDTA. Mitotic cells were collected by shakeoff. All cells were washed once with PBS, and resuspended in TRIzol Reagent. All RNA isolations were performed with Phasemaker™ tubes according to the manufacturer’s protocol (Invitrogen, A33248). Total RNA was resuspended in water with 4 pg spike-in RNA (equal mixture of in vitro transcribed Fluc and Nluc) per 2 ug total RNA for experiments where spike-in RNA was used for normalization. Concentration measurements were carried out using Qubit™ RNA Broad Range Assay Kit (Invitrogen, Q10210).

For HeLa RNA sequencing experiments, rRNA was depleted and libraries were prepared using the KAPA RNA HyperPrep Kit with RiboErase (HMR) (Roche KK8560) according to manufacturer’s instructions. Sequencing was performed on an Illumina NovaSeq SP, in a paired-end 50× 50 mode.

For HCT116 RNA sequencing, total RNA was isolated as described above. rRNA was depleted and libraries prepared using the Watchmaker QiaSeq FastSelect Ribodepletion kit according to manufacturer’s instructions. Sequencing was performed on Element AVITI, in a paired-end 75x75 mode.

For all sequencing experiments reported, poly(A) reads were trimmed using cutadapt (v4.8) with the parameters ‘--minimum-length 1 -a A{25} -A A{25}’. Reads were then mapped to the human genome (Gencode release 25, GRCh38.p7, downloaded from the GENCODE website) using STAR (v2.7.1a; Dobin *et al*, 2013) with the parameters ‘--runThreadN 24 --runMode alignReads --outFilterMultimapNmax 1 -- outFilterType BySJout --outSAMattributes All --outSAMtype BAM SortedByCoordinate’. Exon mapping reads were quantified using htseq-count (v1.99.2; Anders *et al*, 2015) with the parameters ‘-f bam -t exon -r pos -s reverse’. A read cuttoff of ≥25 reads in each sample was applied for each gene.

### Half-life Calculations

mRNA read counts were normalized to spike-in or mitochondrial RNA read counts in pharmacologically transcriptionally inhibited and uninhibited cells, respectively. Normalized abundance values were averaged between two biological replicates and fit to the exponential decay function:

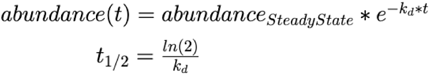

Transcripts with the fit *R*^2^ < 0.8 in interphase cells were excluded from subsequent analysis (Supp. Fig. 2B). Calculated half-life values were arbitrarily capped at 200 h for half-lives >= 200 h, or 0.25 h for half-lives <= 0.25 h for ease of visualization due to their lack of biological interpretability. For the THZ1 experiment (Supp. Fig. 2C), due to technical considerations normalization was done using read counts for top 70 most stable transcripts instead of spike-in read counts, and a fit cutoff of G2 *R*^2^ < 0.4 was used.

### PAL-seq

Sequencing of endogenous mRNA poly(A) tail lengths in interphase and mitosis was performed with PAL-seq v4 as described in (Xiang & Bartel, 2021). The poly(A) tail length was determined using a Hidden Markov Model trained on randomly picked 1% of the filtered read clusters (but no more than 50,000 and no less than 5000) for each library as described in (Xiang *et al*, 2024).

### AID for PABP Depletion

HCT116 cells with PABPC1&4 AID were kindly gifted by the Bartel lab (Xiang & Bartel, 2021). To deplete PABPC1&4 during mitosis, cells were grown to 20% confluency before 8 uM STLC was added together with 1 µg/ml doxycycline to induce OsTIR1 expression and 0.2 mM auxinole (Aobious, AOB8812) in DMSO to prevent premature OsTIR1 activity. An equivalent volume of DMSO was added to non-induced control cells. After 16 h, fresh media with 8 uM STLC was added to all cells, with 0.5mM IAA only added to induced cells. For Western blotting, cells were collected at 0 h, 1 h, or 4 h after IAA addition. For sequencing, cells were collected as described above at 0 h and 4 h. For imaging, SiR-DNA was added (1:10,000), and imaging was started 1 h after addition of IAA and SiR-DNA.

### Western Blotting

Cell pellets were washed once with PBS and resuspended in 1× Laemmli sample buffer (100 mM Tris pH6.8, 12.5% glycerol (v/v), 1% SDS (w/v), 0.1% bromophenol blue (w/v), 200 mM β-mercaptoethanol) then boiled at 95 °C for 5 min. Cell lysates were sonicated at 10% amplitude for 5 s using the Branson Digital Sonifier 450 Cell disrupter to shear genomic DNA. Samples were separated by SDS–PAGE using a 10% gel and transferred to PVDF membrane (VWR). Blots were rinsed once with TBST then blocked in 5% milk at room temperature for 1 h. Blots were then incubated with primary antibodies diluted in 5% overnight at 4 °C. The next day, the blots were washed for 5 min with TBST 4×, incubated with secondary antibody in 5% milk for 1 h at room temperature, followed by 4x more TBST washes and 1x PBS rinse. Blots were imaged using an Odyssey Clx machine (LI-COR) and analysed with the Image Studio software (LI-COR).

Primary antibodies used in this study were anti-PABP1 (1:1,000, Cell Signaling Technology #4992) and anti-alpha tubulin (1:5,000, Sigma-Aldrich, T9026). Secondary antibodies used in this study were IRDye 680RD Goat anti-Mouse (LI-COR 92668070), IRDye 800CW Goat anti-Rabbit (LI-COR 92632211) at 1:10,000 dilution.

### Live-cell Imaging

For live-cell fluorescence imaging, cells were seeded into 12-well glass-bottomed wells (Cellvis, P12-1.5P) and grown to 20% confluence. PABPC1&4 were depleted as described above. DNA was stained with 1:10,000 SiR-DNA. Images were acquired on a Nikon eclipse microscope equipped with a sCMOS camera (ORCA-Fusion BT, Hamamatsu) using a Plan Fluor 20×/0.5 NA objective at 10 min intervals. Time-lapse videos were analyzed using FIJI (ImageJ; NIH). STLC-arrested cells were identified by their characteristic monopolar spindles. Slippage was determined as the first frame in which the individual chromosomes were no longer condensed and timing was calculated using the MTrackJ plugin.

---

**Figure EV1.**
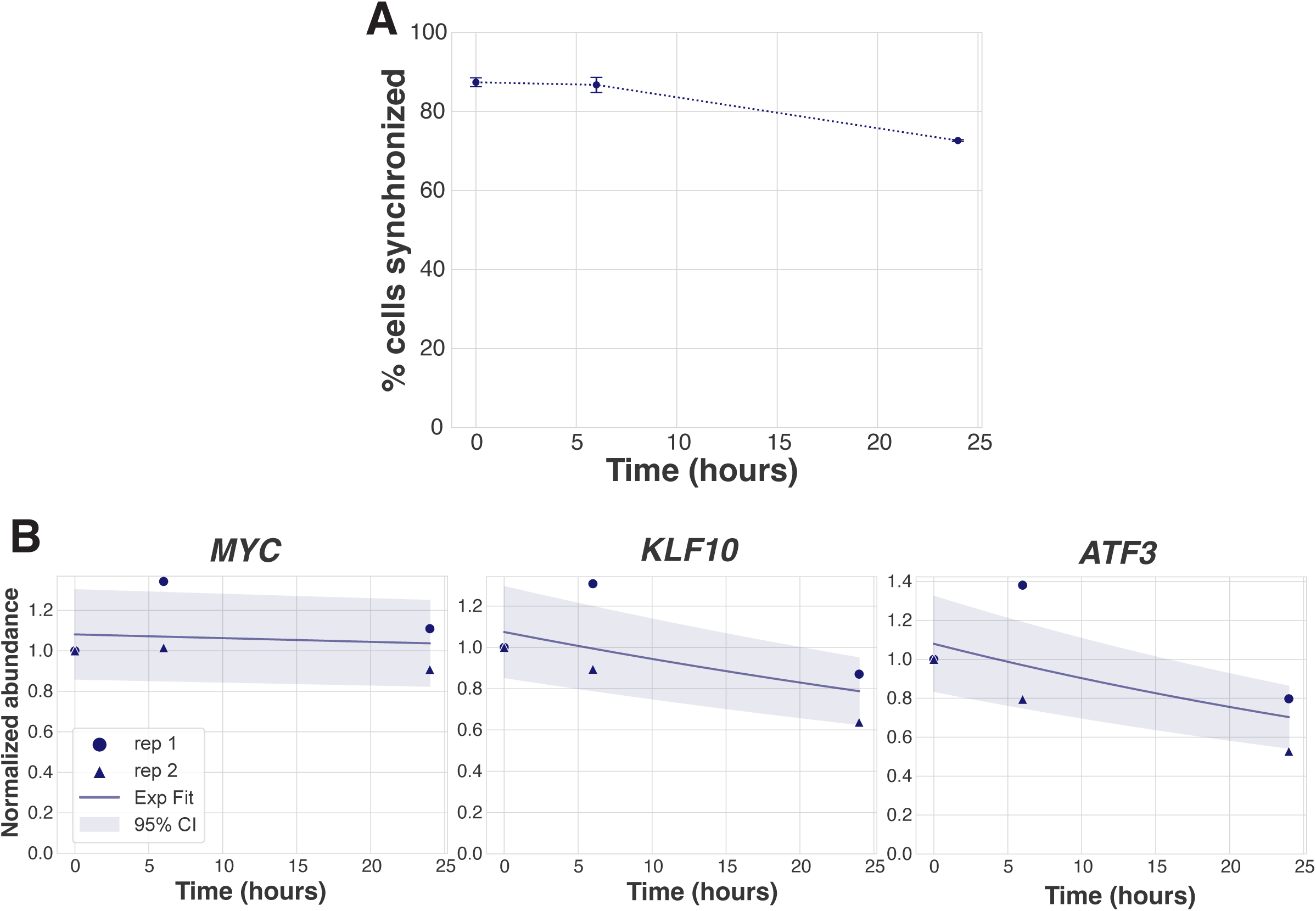
**A)** Percentage of cells synchronized in mitosis during each experimental timepoint (Figure 1), as quantified by FACS analysis of DNA content and pH3(Ser10). **B)** Exponential curve fit to individual transcript abundances in STLC-arrested cells, plotted with 95% confidence intervals. *MYC* t_1/2_ 399 h, *R*^2^ 0.05. *KLF10* t_1/2_ 54 h, *R*^2^ 0.72. *ATF3* t_1/2_ 39 h, *R*^2^ 0.78.

**Figure EV2.**
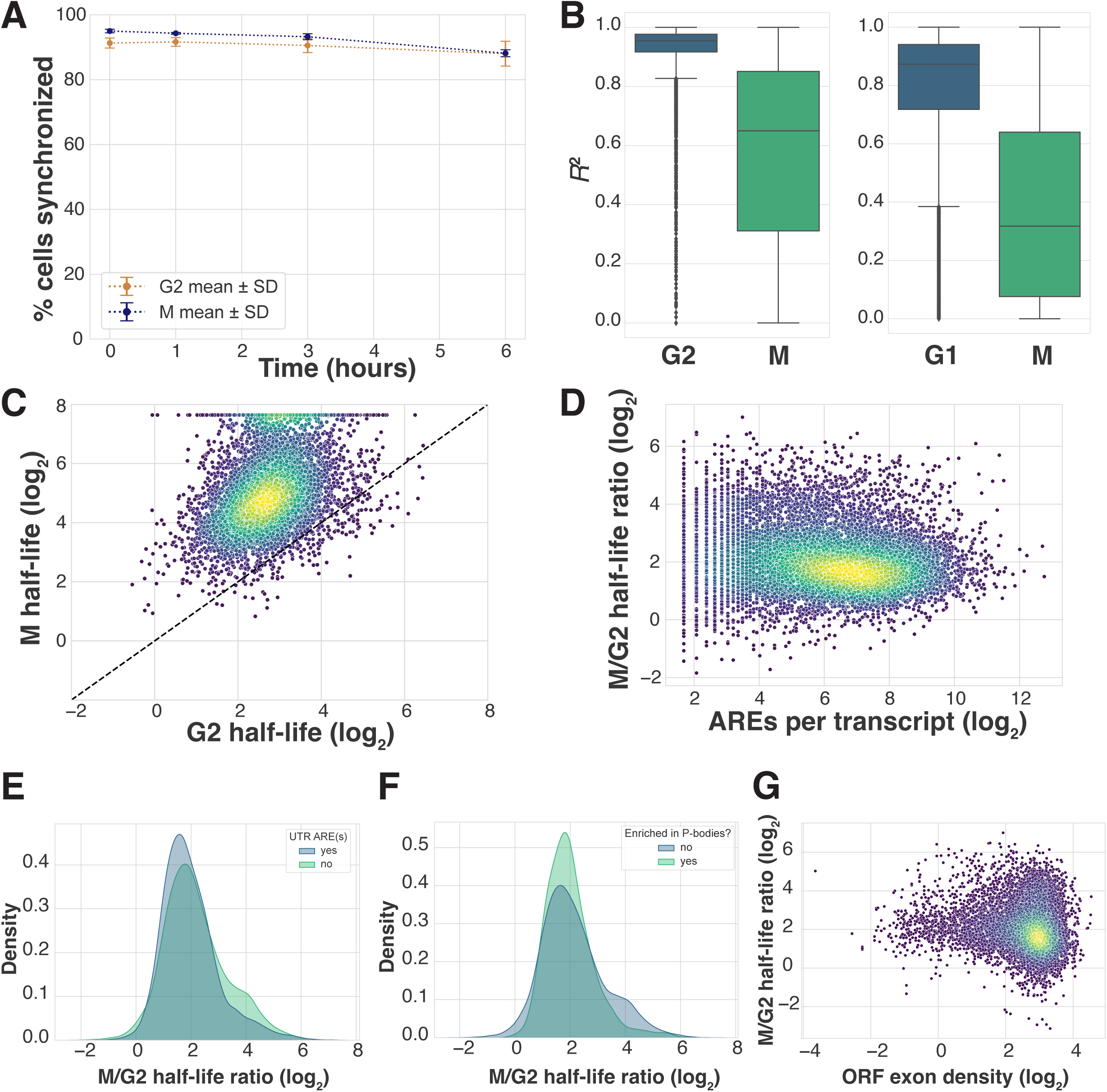
**A)** Percentage of cells synchronized in G2 or mitosis during each experimental timepoint (Figure 2), as quantified by FACS analysis of DNA content and pH3(Ser10). **B)** Boxplots showing *R*^2^ goodness of fit values for half-life calculations for transcripts mapped in transcription inhibition experiments (Fig. 2 B, C). *R*^2^ cutoffs were chosen based on interphase fits in order to allow for the possibility of global mRNA stabilization, as lack of degradation is expected to result in poorer fit values. **C)** Scatterplot comparing mRNA half-lives in cells synchronized in G2 or M and treated with THZ1. Data from one biological replicate. *R*^2^ cutoff > 0.4 was used, n = 6410. Median t_1/2_ 6.77 h and 33.44 h for G2 and M, respectively. **D)** Scatterplot comparing mitotic stabilization and number of AREs per transcript. ARE annotations were acquired from AREsite2 (Fallmann *et al*, 2016). Arbitrary c=0.9 was added to all ARE counts to allow log-transformation of the data. *R_S_* = –0.13. **E)** Probability density function plot comparing degree of mitotic stabilization among transcripts with or without AREs in the 3’ UTR. 3’UTR ARE annotations were acquired from https://brp.kfshrc.edu.sa/ared/Home/FTP (Bakheet *et al*, 2018). n = 3085 transcripts with AREs in 3’ UTR, n = 6773 transcripts with no AREs in the 3’ UTR. **F)** Probability density function plot comparing degree of mitotic stabilization among transcripts based on their P-body enrichment status (Hubstenberger *et al*, 2017). P-body enrichment was defined by FDR < 0.01 and log_2_ enrichment > 2, resulting in 1602 P-body-enriched transcripts and 8256 not enriched. **G)** Scatterplot comparing degree of mitotic stabilization and ORF exon density, as calculated in (Agarwal & Kelley, 2022). *R_S_*= –0.15.

**Figure EV3.**
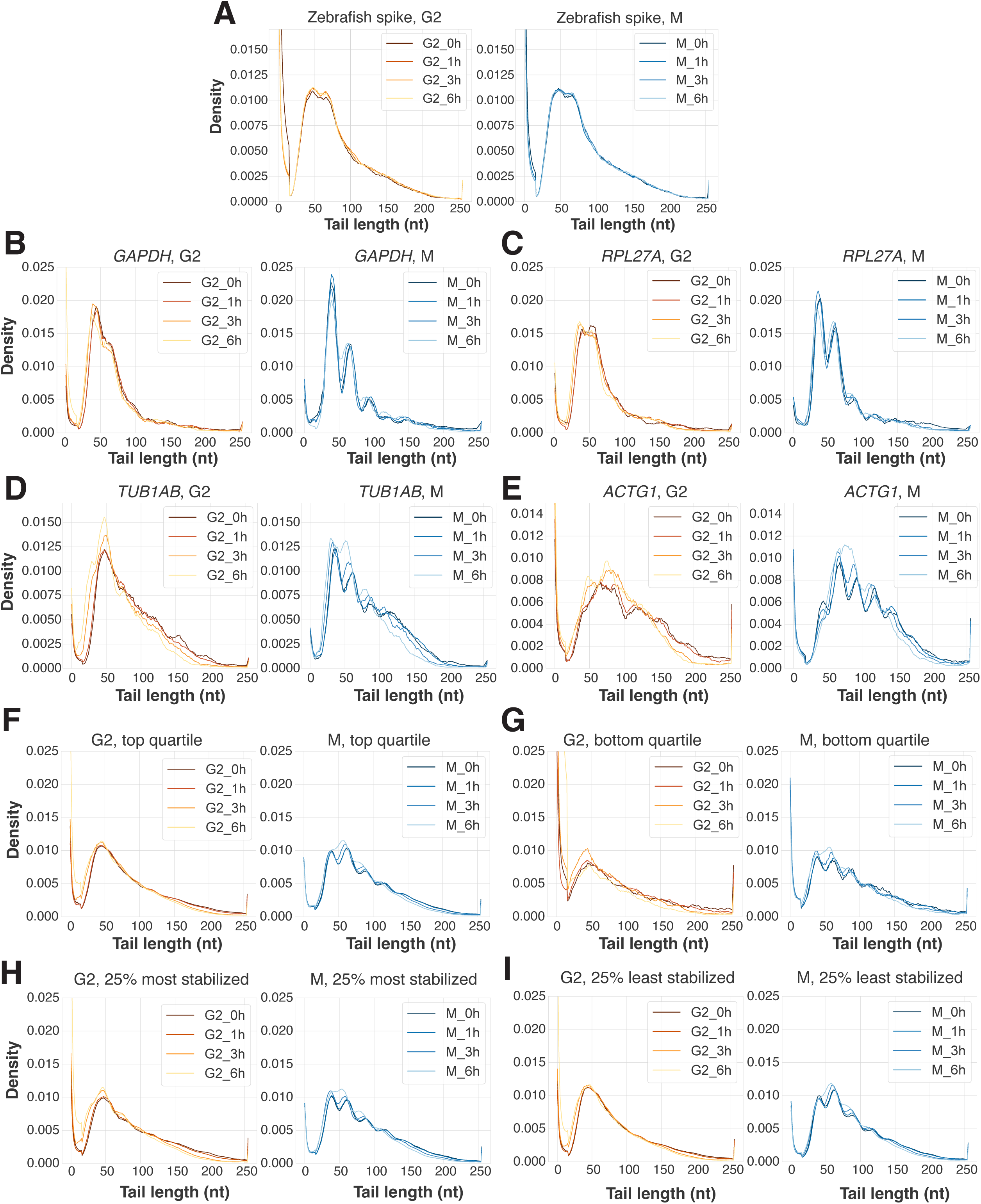
**A)** Poly(A) tail length distributions from zebrafish mRNA spiked into G2- or M-treated samples as measured by PAL-seq. **B–E**) Poly(A) tail length distributions in G2- or M-arrested cells treated with actinomycin D as measured by PAL-seq for GAPDH (**B**), RPL27A (**C**), TUB1AB (**D**), ACTG1 (**E**). **F–G**) Poly(A) tail length distributions in RO-3306- or STLC-arrested cells treated with actinomycin D as measured by PAL-seq for 25% most highly expressed genes (**F**) or 25% mostly lowly expressed genes (**G**). **H–I**) Poly(A) tail length distributions in G2- or M-arrested cells treated with actinomycin D as measured by PAL-seq for 25% most stabilized transcripts (**H**) or 25% least stabilized transcripts (**I**) based on analysis from Figure 2.

**Figure EV4.**
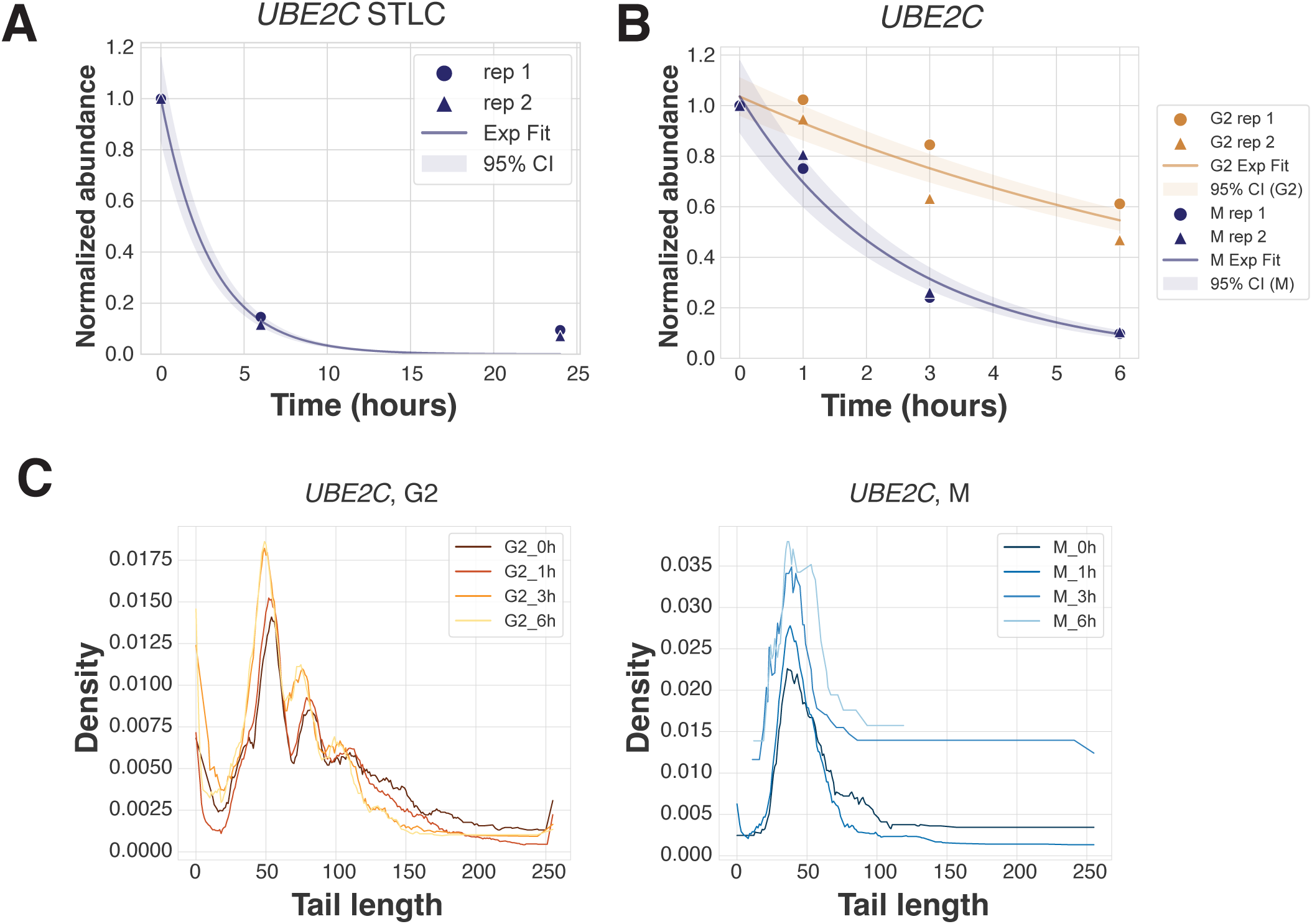
**A)** Exponential curve fit to *UBE2C* transcript abundances over 24 hours of mitotic arrest by STLC, plotted with 95% confidence intervals. t_1/2_ 2.05 h, *R*^2^ 0.99. **B)** Exponential curve fit to *UBE2C* transcript abundances over 6 hours of G2 or mitotic arrest with transcription inhibition by actinomycin D, plotted with 95% confidence intervals. t_1/2_ 6.27 h and 1.75 h, *R*^2^ 0.97 and 0.98 in G2 and M, respectively. **C)** Poly(A) tail length distributions in G2- or M-arrested cells treated with actinomycin D as measured by PAL-seq for *UBE2C*.

**Figure EV5.**
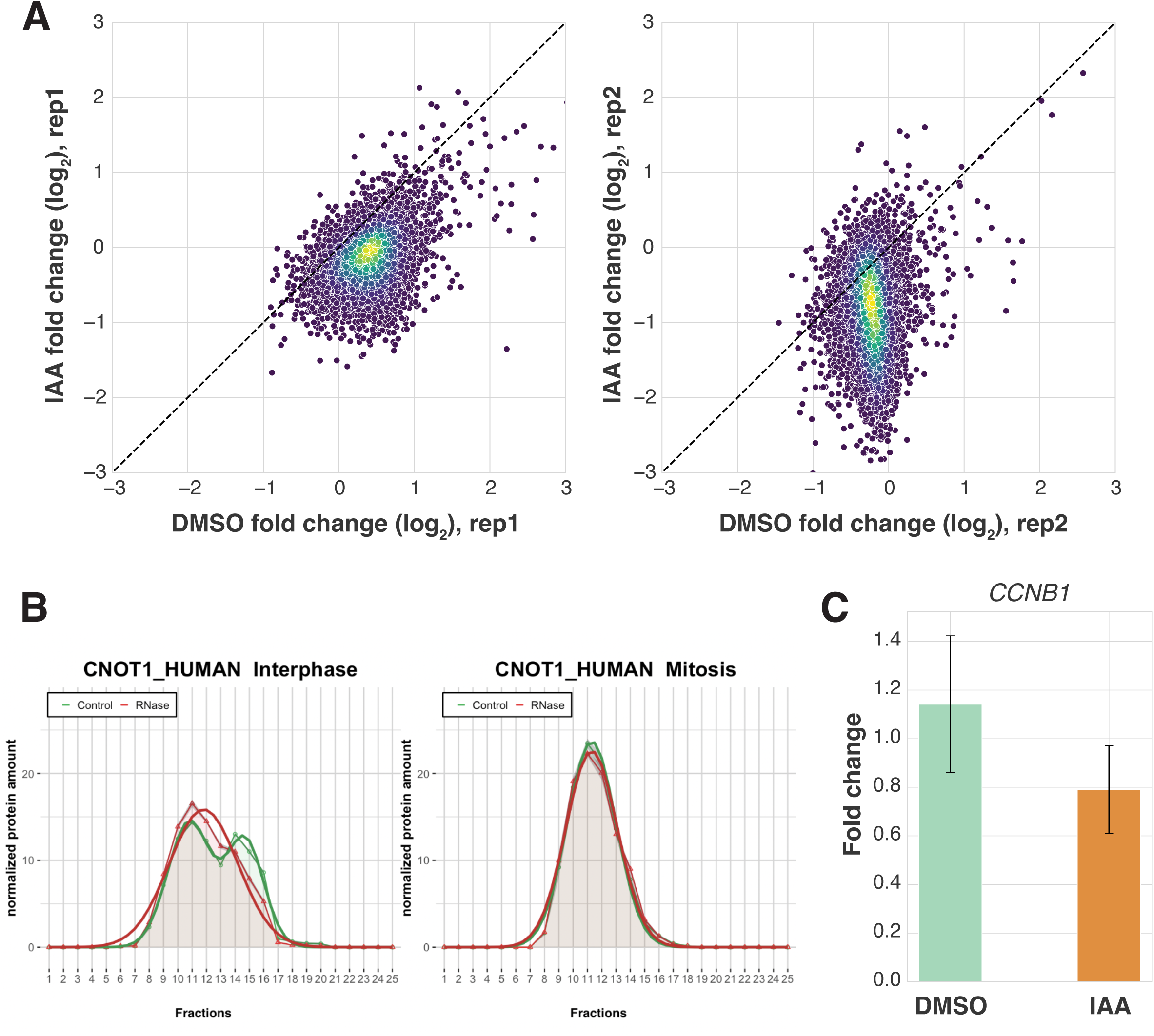
**A)** Scatterplots comparing mRNA abundance fold changes in STLC-arrested HCT116 cells within 4 hours of either DMSO control treatment or IAA-induced PABPC1&4 depletion. Replicates 1 & 2, n = 11078. **B)** Graphical representation of the CNOT1 protein amount from interphase or mitotic HeLa cells in 25 different fractions of control (green) and RNase-treated (red) sucrose density gradients analyzed by mass spectrometry. Plots were acquired from https://r-deep3.dkfz.de/ (Rajagopal *et al*, 2025). The leftward shift of the distribution upon RNase treatment in interphase cells suggests interaction with RNA, whereas a lack of shift in mitotic cells suggests reduced or no interaction. **C)** Bar graph comparing *CCNB1* degradation in STLC-arrested HCT116 cells within 4 hours of either DMSO control treatment or IAA-induced PABPC1&4 depletion. n = 2 biological replicates.

